# ZNF191 Inhibits Hepatocellular Carcinoma Metastasis via DLG1-mediated YAP1 Inactivation

**DOI:** 10.1101/027060

**Authors:** Di Wu, Haitao Li, Yufeng Liu, Hexige Saiyin, Chenji Wang, Zhen Wei, Wenjiao Zen, Danyang Liu, Qi Chen, Haojie Huang, Guoyuan Liu, Songmin Jiang, Long Yu

## Abstract

Searching targets for hepatocellular carcinoma (HCC) treatment, we identified zinc finger protein 191 (ZNF191) as a suppressor against HCC metastasis. Over-expressing ZNF191 in HCC cells impaired cell motility, while ZNF191 depletion promoted HCC cell migration *in vitro* and metastasis *in vivo* through triggering yes-associated protein 1 (YAP1) signaling. Chromatin immunoprecipitation-sequencing (ChIP-seq) revealed that ZNF191 specifically bound to the promoter of *Discs, Large homolog 1* (*DLG1*), a cell polarity maintainer and a negative regulator of YAP1. Double-knockdown experiments showed that DLG1 was not only the mediator of ZNF191’s function to suppress migration but also a link between ZNF191 and YAP1 signaling. ZNF191 was down-regulated in metastatic HCCs, correlating positively with DLG1 levels and inversely with YAP1 activation. Our findings indicate ZNF191 functions as a metastasis suppressor via DLG1-mediated YAP1 signaling inactivation.

## List of Abbreviations

ChIP-seq: Chromatin immunoprecipitation-sequencing
DLG1, Discs,: large homolog 1
EMSA: Electrophoretic mobility shift assay
HCC: Hepatocellular carcinoma
mRNA: messenger RNA
qPCR: Real-time quantitative polymerase reaction
RNAi: RNA interference
shRNA: short hairpin RNA
siRNA: small interfering RNA
WNT: Wingless-type
YAP1: Yes-associated protein 1
ZNF191: Zinc finger protein 191.

## Introduction

Much due to the propensity for metastasis, HCC remains one of the leading fatal cancers^1^. Identifying prognostic markers and treatment targets for metastatic HCC is in urgent need.

With the complex progression of solid tumors, cancer cells gradually delaminated and intravasated^2^, both associating with loss of cell polarity^3^. The SCRIB-LGL-DLG complex is a conserved module in maintaining apical-basolateral polarity in both Drosophila and mammal cells^44^,^5^. All its components are tumor suppressors in *Drosophila*^6^ and predominantly inhibit cancer cell migration and metastasis^4^,^7^. Nevertheless, how this complex is deregulated during tumor progression is scarcely known.

SCRIB-LGL-DLG complex crosstalks with the Hippo-YAP signaling pathway which takes important part in HCC progression^4^. Accumulating clues strongly implicate the role of SCRIB-LGL-DLG complex as a negative regulator of YAP/TAZ signaling^8^-^11^.

ZNF191 (also called ZNF24) was previously identified as a proliferation promoter of HCC cells^12^. More than a transcription factor specifically recognizing TCAT repeating motifs^13^, it was also enriched at nascent DNA and facilitating DNA replication^14^. ZNF191 was even reported to facilitate the migration of endothelial cells^15^ and vascular smooth muscle cells^16^. However, we further found that ZNF191 should be lost in metastatic HCCs. We have confirmed that ZNF191 is indeed a suppressor against cell migration and metastasis, meanwhile an inhibitor of YAP1. ZNF191 is a direct and specific transcriptional regulator of DLG1. The unexpected function of ZNF191 is mediated by a putative DLG1-YAP1 signaling.

## Results

### Down-regulation of ZNF191 associates with metastasis and poor clinical outcomes in HCC

We previously found ZNF191 was up-regulated in a subset of HCC specimens, promoting the proliferation of non-migratory L02 and Hep3B cells *in vitro* and *in vivo*^12^.

To further investigate the role of ZNF191in HCC progression, we collected a cohort of 120 HCC specimens with or without intrahepatic metastasis. We first identified a ZNF191 antibody that is suitable for Immunohistochemistry (IHC) (Supplementary Fig. 1) and then applied this antibody to detect ZNF191 expression in these samples. By examining the correlation between ZNF191 expression and clinicopathological features (Supplementary Table 1), we found ZNF191 was over-expressed in 73% HCC cases without metastasis but almost lost in 64% metastasis-present cases (Fig. 1a, b). The inverse correlation was statistically significant (*P* < 0.01).

**Figure 1.**
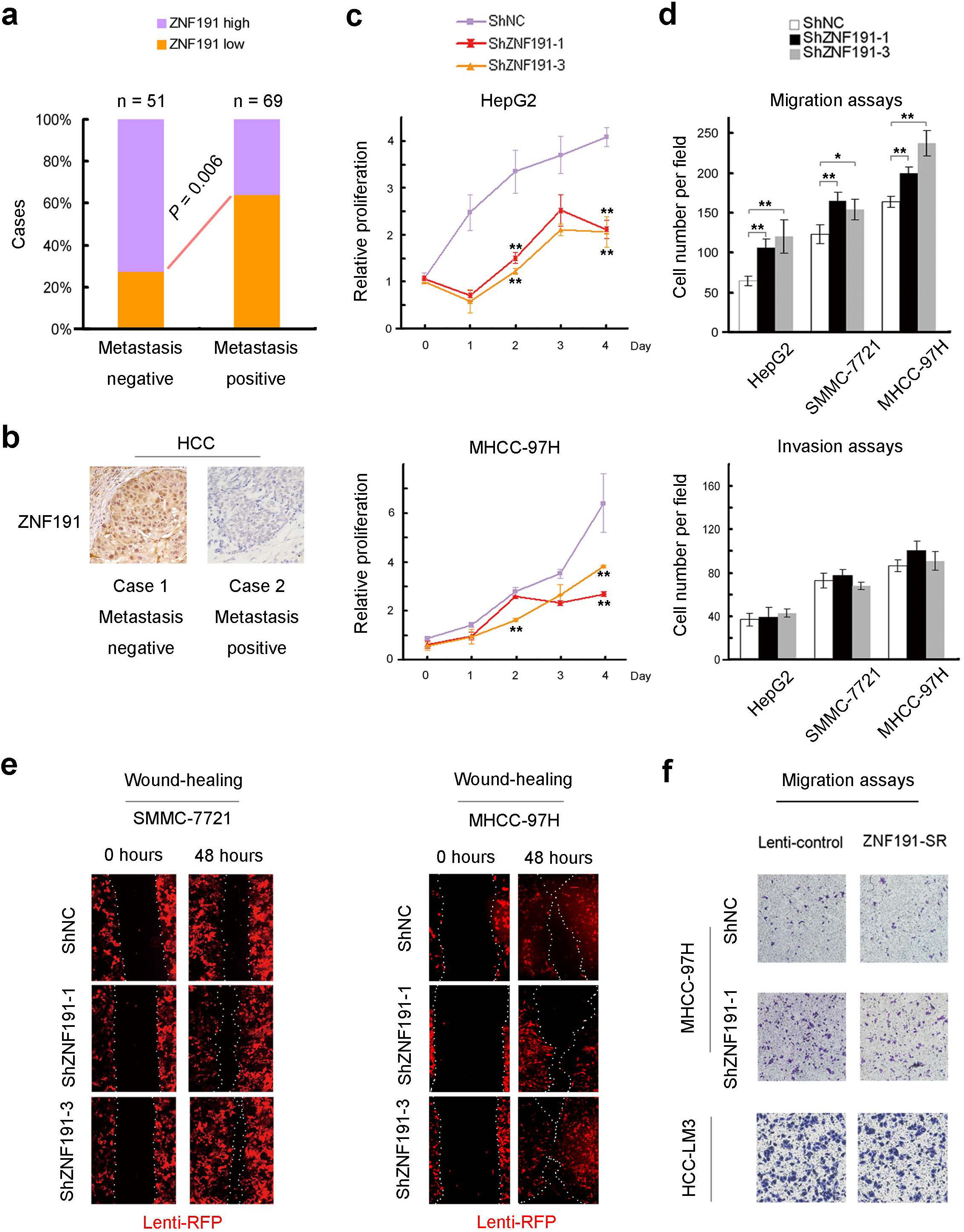
ZNF191 was down-regulated in metastatic HCCs and suppressed HCC cell migration *in vitro*. (a) ZNF191 expression levels in cancerous tissues were scored as higher or lower compared to non-cancerous liver tissues (also see Supplementary Fig. 1c). (b) Representative ZNF191 staining in metastasis-negative/positive HCC. (c) Proliferation assays using MTS method were performed. (d) - (f) HCC cells transfected with lentivirus expressing short hairpin RNAs (shRNAs) or cDNA had been maintained by puromycin selection for at least 2 weeks before experiments. The efficacy of ZNF191 depletion was validated in Fig. 3a, b. The efficacy of ZNF191 over-expressing or rescuing was validated in Supplementary Fig. 9a-e. (d) The *in vitro* migration and invasion potentials of 3 lines of HCC cells stably transfected with negative control (NC) or ZNF191-depleting shRNAs were assayed and analyzed (also see Supplementary Fig. 4d, e). (e) Would-healing assays were applied to measure the mixed effects of cell growth and migration. Red fluorescence protein (RFP) is the tracing mark of these lentivirus transfected stable lines. (f) Over-expressing ZNF191 in MHCC-97H and HCC-LM3 cells, or rescuing ZNF191 expression in ZNF191-knockdown MHCC-97H cells was performed using a lentivirus expressing a synonymously mutated ZNF191 CDS that is insensitive to ShZNF191-1 (ZNF191-SR), for details see **Online Methods**. These cells were assayed for their migration abilities and representative results are shown. Statistic analysis of the results is in Supplementary Fig. 5a, b. * *P* < 0.05; ** *P* < 0.01.

Accordingly, in the published databases we also found ZNF191 was over-expressed mainly in metastasis-free HCCs but down-regulated in HCCs with intrahepatic metastasis, distal metastasis (not significantly) or macroscopic vascular invasion (Supplementary Fig. 2a-d). Moreover, Kaplan-Meier analyses in TCGA database demonstrated that ZNF191 low expression predicts significantly shorter overall survival (Supplementary Fig. 2e, f), further suggesting a potential use of ZNF191 as a favorable prognostic marker in HCC patients.

### ZNF191 suppresses HCC cell migration *in vitro* and metastasis *in vivo*

Reduced or lost expression of ZNF191 in metastatic HCCs prompted us to investigate the role of ZNF191 in cell migration and invasion. To this end, we depleted ZNF191 in an array of HCC cell lines to test their migration ability. To our surprise, ZNF191 knockdown inhibited cell proliferation (Fig. 1c), but stimulated most of these HCC cell lines to migrate (Fig. 1d & Supplementary Fig. 4c-e).

In wound-healing assays, the mixed effect of cell growth and migration was tested. ZNF191-knockdown SMMC-7721 cells recovered faster than control cells (Fig. 1e). Similarly, ZNF191 knockdown in MHCC-97H cells increased migration so that they should recover equally or faster than the control cells, considering ZNF191 knockdown had decreased growth in these cells (Fig. 1c the panel below). We ectopically expressed ZNF191 using lentivirus in highly migratory HCC-LM3 cells (which express over 10 fold less ZNF191 than HepG2 cells, see Supplementary Fig. 4a, b) and ZNF191-knockdown MHCC-97H cells. All these treatments impaired cell motility (Fig. 1f). However, ZNF191 knockdown had no overt effect on the invasiveness of HCC cells (Fig. 1d the panel below).

To further clarify the role of ZNF191 in HCC metastasis. ZNF191-knockdown MHCC-97H cells were orthotopically implanted into the livers of nude mice. 4 weeks and 8 weeks after implantation, *in vivo* imaging indicated that ZNF191 knockdown tumor-bearing mice suffered from smaller orthotopic tumors at first (Fig. 2a) but finally showed significantly more whole body spreading and more metastatic nodules in lungs than mice bearing control tumors (Fig. 2a, d), which was confirmed by tracing exogenous RFP marker under confocal microscope and histological examination (Fig.2b, c). We further found ZNF191 knockdown resulted in more circulating tumor cells in mouse blood (Fig. 2e). These findings indicate that ZNF191 inhibits HCC metastasis via suppressing primary tumor cells migration and intravasation.

**Figure 2.**
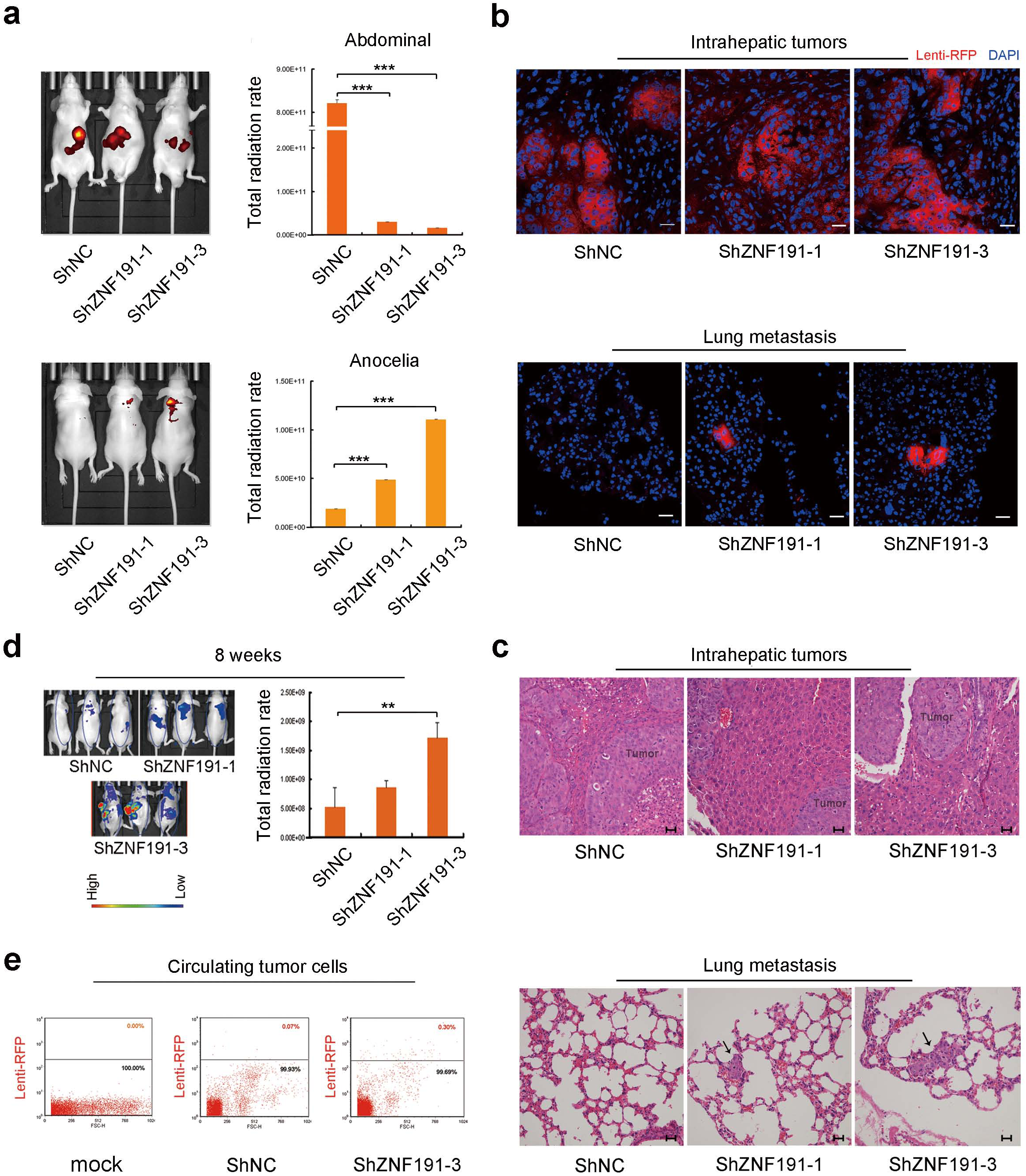
ZNF191 inhibits HCC cell metastasis. Different MHCC-97H stable lines were orthotopically implanted into the livers of nude mice. For each group (n = 8), 4 weeks and 8 weeks after transplantation, all the mice were anaesthetized and imaged. Exogenous cancer cells had been labeled with RFP by lentivirus. (a) Left column: the signal in the ventral view (the upper panel) mainly represented the orthotopic tumors and their abdominal metastasis; and the dorsal view (the panel below) showed the signal from anocelia metastasis mainly in lungs. (a) Right column: The statistic analysis of the results in the left column, quantification of the fluorescence signal was performed using the animal *in vivo* imaging system and software. (b) For each group, a portion of the animals (n = 3) were sacrificed and anatomized. Livers and lungs of the mice were embedded with OTC compound and frozen, and then sliced as 5μm thick sections to be observed under confocal microscope. (c) Lungs of the mice were fixed and paraffin-embedded. 4μm thick sections were sliced. Hematoxylin-eosin (H&E) staining results of these sections were presented. Arrows show the metastasis sites in lungs. (d) 8 weeks after transplantation, the surviving mice of each group were anaesthetized and imaged (2 mice in ShZNF191-1 group had died from too heavy tumor loads in this period). The fluorescence signal in the same size of body region was quantified. Statistic analysis of the quantification results is shown. The ventral view could not be achieved because the tumors had grown too large. (e) The circulating tumor cells (CTCs) in the peripheral blood of tumor-bearing mice were counted using flow-cytometry detecting RFP signal. For each counting, an equal number of 10,000 cells were counted as the loading control. The threshold lines in the panels were set arbitrarily, in purpose to discern differences between groups. Statistic analysis is shown in Supplementary Fig. 5c. ** *P* < 0.01; *** P < 0.001; Bar = 25 μm.

### ZNF191 suppresses cell migration independently of E-cadherin, β-catenin or VEGF

To explore the mechanisms underlying the metastasis-suppressive function of ZNF191, we first sought to determine whether epithelial-mesenchymal transition (EMT) had occurred after ZNF191 knockdown. We found that neither expression of E-cadherin nor N-cadherin was changed after ZNF191 knockdown (Fig. 3a, b) Moreover, in our SMMC-7721 cells which lack expression of E-cadherin, ZNF191 knockdown still converted the cells from the round to the spindle-like shape and increased expression of Vimentin (Supplementary Fig. 3a-c), suggesting that the effect of ZNF191 on cell migration is independent of E-cadherin.

**Figure 3.**
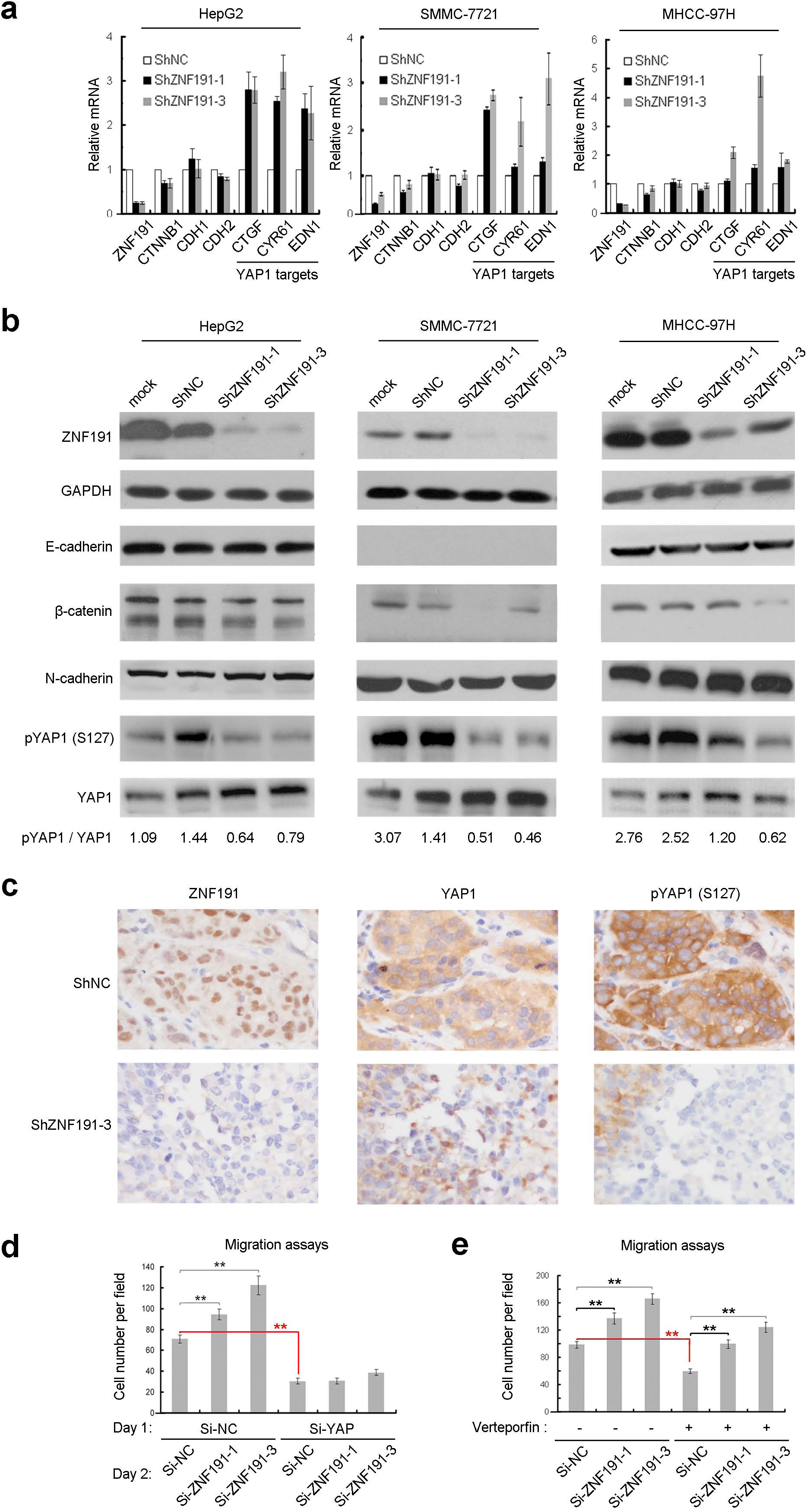
Depleting ZNF191 in HCC cells resulted in the activation of YAP1 signaling. (a) Fold changes of mRNA levels were quantified by qPCR in 3 HCC cell lines. (b) Equal amount of cell lysates were subjected to Western blotting. GAPDH was detected as the loading control. The two-band blotting of `β`-catenin is detailed in **Cell line identification and choosing** section of **Online Methods**. Grayscale of each band was quantified via Quantity One software as signal intensity. The ratio of the signal intensity of phosphorylated YAP1 (pYAP1) to that of total YAP1 was designated as the inactivation level of YAP1 signaling. (c) Paraffin-embedded consecutive sections of orthotopic tumors in mice livers were subjected to IHC staining. (d) Cells were pre-depleted of YAP1 by siRNA for 24 hours and then transfected with ZNF191-depleting siRNAs for 24 hours. Then treated cells were subjected to migration assays. The efficacy of YAP1 knockdown is shown in Supplementary Fig. 6h. (e) During the RNAi and migration assays, Verteporfin final concentration in the medium was 10 μM (DMSO as negative control); the experiment was carried out in darkness. ** *P* < 0.01.

`β`-catenin down-regulation was moderate in these cells after ZNF191 knockdown (Fig. 3a, b) and impossible to mediate the migration increase, because we confirmed that WNT3A or `β`-catenin would promote rather than inhibit the migration of HepG2 cells (Supplementary Fig. 6a, b). ZNF191 was reported to suppress VEGF^17^,^18^, but ZNF191 depletion still increased cell migration to the same fold when VEGF was blocked by Bevacizumab (AVASTIN), an inhibitor of VEGF-A (Supplementary Fig. 6c).

### ZNF191 knockdown results in activation of YAP1 signaling

Incidentally, mining transcript profiling data in HEK293 cells^19^, we found that many well-studied target genes of the transcriptional co-activator YAP1 (*CTGF*, *CYR61*, *EDN1* etc.) also happen to be the targets of ZNF191 (Supplementary Fig. 7a). More intriguingly, in a database of 883 lines of cancer cells, ZNF191 expression is inversely correlated with almost all the prominent co-target genes of YAP1, including *CTGF*, *CYR61* and *EDN1* (Supplementary Fig. 7d, e). Indeed, our microarray data indicated that *CTGF* and *CYR61* were significantly up-regulated upon ZNF191 knockdown in L02 cells (Supplementary Fig. 7f).

Considering the pro-metastasis role of YAP1^20^,^21^, we repeated the experiments and confirmed that YAP1 target genes were up-regulated after ZNF191 stable knockdown both in HEK293 (Supplementary Fig. 7c) and in HCC cell lines (Fig. 3a). These findings prompted us to determine whether YAP1 might be activated after ZNF191 knockdown. We demonstrated that phosphorylation of YAP1 at serine-127 was reduced after ZNF191 knockdown (Fig. 3b). In ZNF191 knockdown orthotopic tumors, consecutive sections showed more nuclear staining of YAP1, a hallmark of YAP1 activation, and less pYAP1 staining (Fig. 3c).

To determine whether ZNF191 suppresses metastasis through regulating YAP1, YAP1 signaling was blocked either by knocking down YAP1 or treating HepG2 cell with Verteporfin, a compound that disrupts YAP-TEAD complex^22^. In both cases, migration of HepG2 cells was inhibited (Fig. 3d, e); in YAP1-knockdown cells, depleting ZNF191 no longer stimulated migration (Fig. 3d). These data suggest that YAP1 activation takes an essential part in stimulating cell migration after ZNF191 knockdown.

### DLG1 is the only candidate YAP1 regulator with its promoter directly targeted by ZNF191

Next we sought to determine how the YAP1 signaling is regulated by ZNF191. To genome-widely identify the binding sites of ZNF191, chromatin immunoprecipitation-sequencing (ChIP-seq) was performed using a validated ZNF191 antibody (Supplementary Fig. 8a) in both HEK293 (where ZNF191 regulates YAP1 significantly) and HepG2 cells (a representative of HCC cells). After peak calling, we identified 1315 ZNF191-bound peaks in HepG2 and 646 in HEK293 cells. In these peaks, we identified the most frequent motifs (Fig. 4a). In HEK293 cells, the core motif TCAT triplicates match perfectly with the ZNF191 binding sites defined previously^13^; while in HepG2 cells the core motif TSAW repeats (S = G or C; W = A or T) extend the known ZNF191 motif by limited tolerance in the second and fourth positions. These findings in turn qualified our ChIP-seq results.

**Figure 4.**
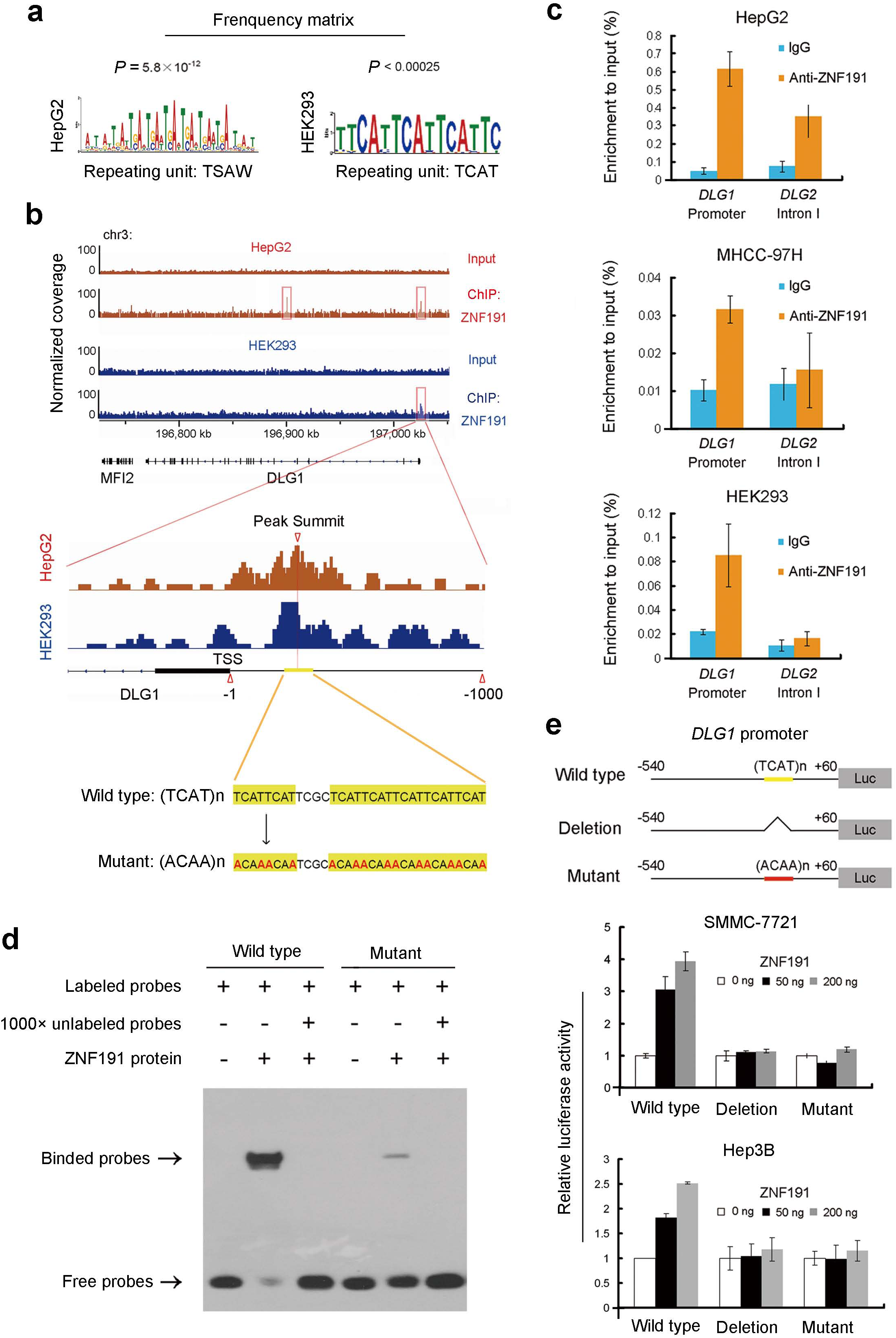
ZNF191 directly binds to the TCAT repeating motifs in *DLG1* promoter and activates it. (a) The top motifs identified by MEME analysis of ChIP-seq-enriched genomic regions in HEK293 and HepG2 cells. (b) Overview of the chromatin immunoprecipitation and high-throughput sequencing (ChIP-seq) results around *DLG1* gene. Peaks were overlaid by sequencing reads. Compared to genome DNA inputs, the DNA regions enriched by ZNF191 immunoprecipitation was identified as ZNF191-bound peaks. The most over-lapped base pair was defined as the peak summit. (The panel below) Demonstration of the symbols and sequence changes in the mutant sequences/probes that will appear in (d) & (e). (c) ChIP-qPCR was performed. Apart from *DLG1* promoter specific primers, primers specific to a region in *DLG2* first intron (another ZNF191-bound peak in HepG2, see Supplementary Fig. 8e) was also applied. (d) ZNF191 recombinant protein was incubated with indicated probes for 20 minutes and subjected to electrophoretic mobility shift assay (EMSA). (e) A portion of *DLG1* promoter (wild type or deletion/mutant) was cloned to pGL3-luciferase vector, which was transfected with pCMV-Myc-ZNF191 and pRL-SV40 vectors into SMMC-7721 or Hep3B cells. 20 hours after transfection, cells were harvested and subjected to dual luciferase assays.

235 peaks in HepG2 and 141 peaks in HEK293 cells locate in the 2,000 basepair regions up-stream of transcriptional start sites (TSSs) (Supplementary Fig. 8b). However, ZNF191-bound peaks were associated with neither YAP/TAZ signaling core elements (*MST1/2*, *LATS1/2*, *YAP1/WWTR1*, *TEAD1/2/3/4*) nor its down-stream target genes (*CTGF*, *CYR61*, *EDN1* etc.).

*DLG1*, a component gene of the cell polarity module SCRIB-LGL-DLG, is an indirect negative regulator of YAP1 (Supplementary Fig. 8c, d). We noticed that in both HEK293 and HepG2 cells, ZNF191 bound to the proximal promoter of *DLG1*. Although ZNF191 bound to other *DLG* family genes at their gene bodies in HepG2 (Supplementary Fig. 8e), *DLG1* is the **only** known YAP1 regulator of which the promoter is targeted by ZNF191 in both cell lines.

In accordance with what we found, just near the summits of the ZNF191-bound peaks upstream *DLG1* in both cells, we found a typical TCAT repeats (7 repeats in total, segmented by a TCGC, see Fig. 4b & Supplementary Fig. 8f). ChIP-qPCR confirmed that endogenous ZNF191 enriched around this motif (Fig. 4c). Electrophoretic mobility shift assay (EMSA) further indicated that recombinant ZNF191 protein directly bound to this sequence with high affinity.

When the repeating unit TCAT was mutated to ACAA, the binding affinity with ZNF191 almost disappeared (Fig. 4d). Accordingly, only the wild type *DLG1* promoter was found to be activated by ZNF191 and when the TCAT repeats were deleted or mutated to ACAA repeats, ZNF191 failed to activate the reporter system in either SMMC-7721 or Hep3B cells (Fig. 4e).

In HCC cells, ZNF191 knockdown resulted in a decrease of *DLG1* expression while the other *DLG* family genes were largely not affected (Fig. 5a, b). And the reduction of *DLG1* was confirmed in ZNF191 knockdown orthotopic tumors previously described (Fig. 5c). These findings indicate that ZNF191 directly activates the *DLG1* promoter and maintained DLG1 expression in HCC cells.

**Figure 5.**
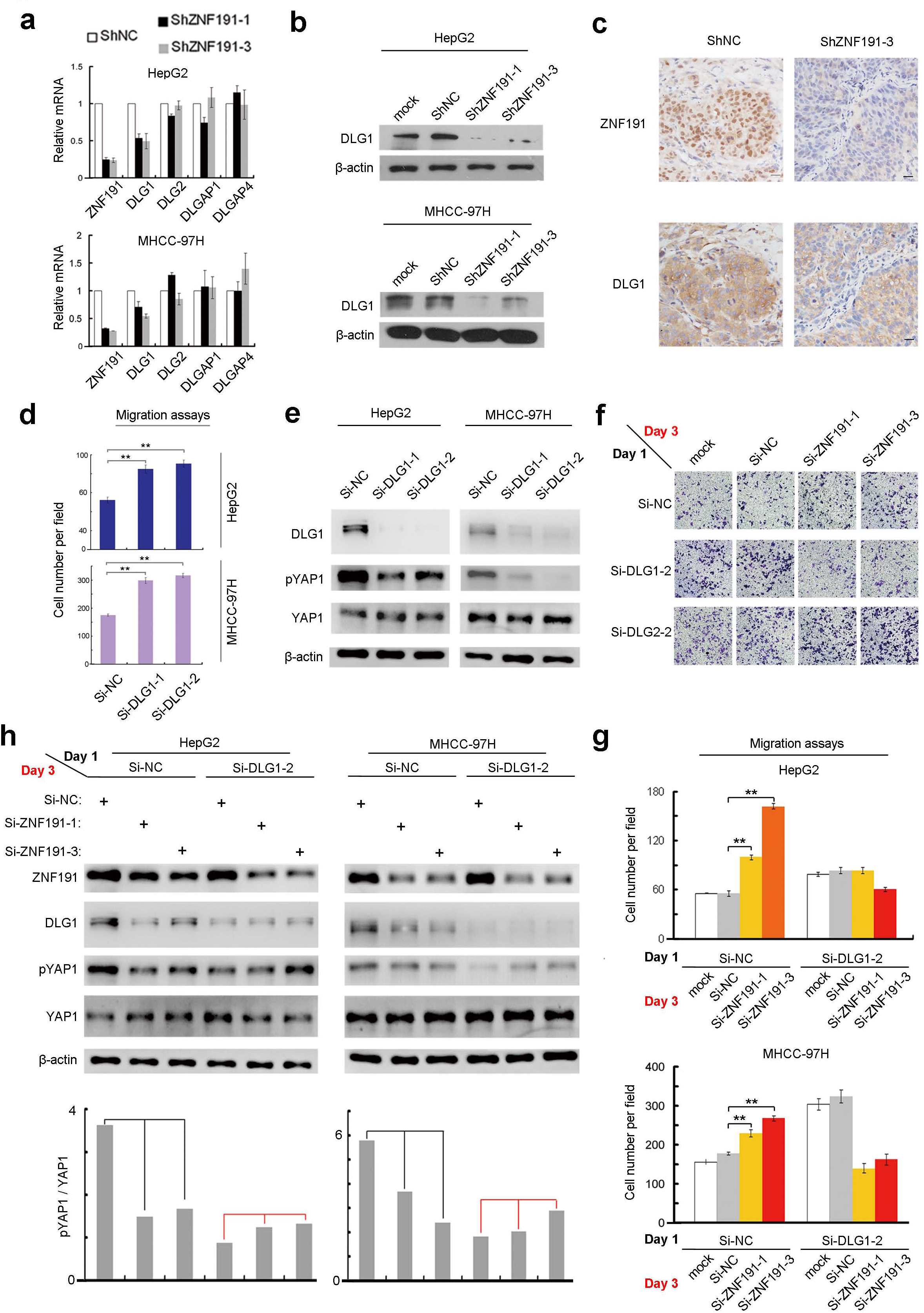
DLG1 is a functional mediator of ZNF191. (a) The same cDNA samples as in Figure 3a were subjected to qPCR. (b) The same cell lysates as in Figure 3b were subjected to Western blotting. `β`-actin was detected as the loading control. (c) Paraffin-embedded consecutive sections of orthotopic tumors in mice livers were subjected to IHC staining. Bar = 25 μm. (d) Cells transfected with siRNAs for 72 hours were subjected to migration assays or (e) to Western blotting. (f) MHCC-97H cells were pre-depleted of DLG1 or DLG2 by siRNAs for 48 hours and then transfected with ZNF191-depleting siRNAs for 24 hours. Treated cells were subjected then to migration assays with the statistic analysis shown in (g). (h) Equal amount of cell lysates were subjected to Western blotting. Grayscale scanning and analysis were performed as in Figure 3b. * *P* < 0.05; ** *P* < 0.01.

### DLG1 inhibits HCC cell migration and mediates the function of ZNF191

Firstly, DLG1 was confirmed to be a migration-suppressor in HCC cells via RNAi (Fig. 5d). DLG1 knockdown resulted in the reduction of YAP1 phosphorylation (S127) (Fig. 5e). Moreover, DLG1 knockdown partially compensated for the migration decrease caused by ZNF191 over-expression or rescue (Supplementary Fig. 9f). In DLG1-depleted MHCC-97H cells, ZNF191 knockdown was no longer able to increase cell migration as it did in control cells (Fig. 5f, g). The similar result was observed in HepG2 cells (Fig. 5g), while DLG2 knockdown did not affect the migration-suppressive potential of ZNF191 (Fig. 5f the bottom panels). These indicate that migration-suppressive function of ZNF191 is DLG1 dependent. Notably, in DLG1-knockdown cells, depleting ZNF191 no longer decreased YAP1 phosphorylation (Fig. 5h). These indicate that DLG1 is down-stream of ZNF191 and up-stream of YAP1.

### ZNF191 expression correlates with DLG1 levels and inversely correlates with YAP1 activation in HCC

To validate our findings, we correlated among expression of ZNF191, DLG1 and YAP1 proteins using IHC in consecutive sections in a cohort of 120 HCC specimens. Nucleus YAP1 was found significantly correlated with intrahepatic metastasis (Fig. 6a), and ZNF191 low expressing cases exhibited more nuclear staining of YAP1 (Fig. 6c). These results support our model that ZNF191 loss induces metastasis through activating YAP1. Staining patterns of ZNF191 and DLG1 were similar (Fig. 6d); their mRNA levels in another 44 HCC samples correlated to each other significantly (Fig. 6e), which is further supported by TCGA data (Fig. 6f). These observations support our model that ZNF191 maintains DLG1 transcription.

**Figure 6.**
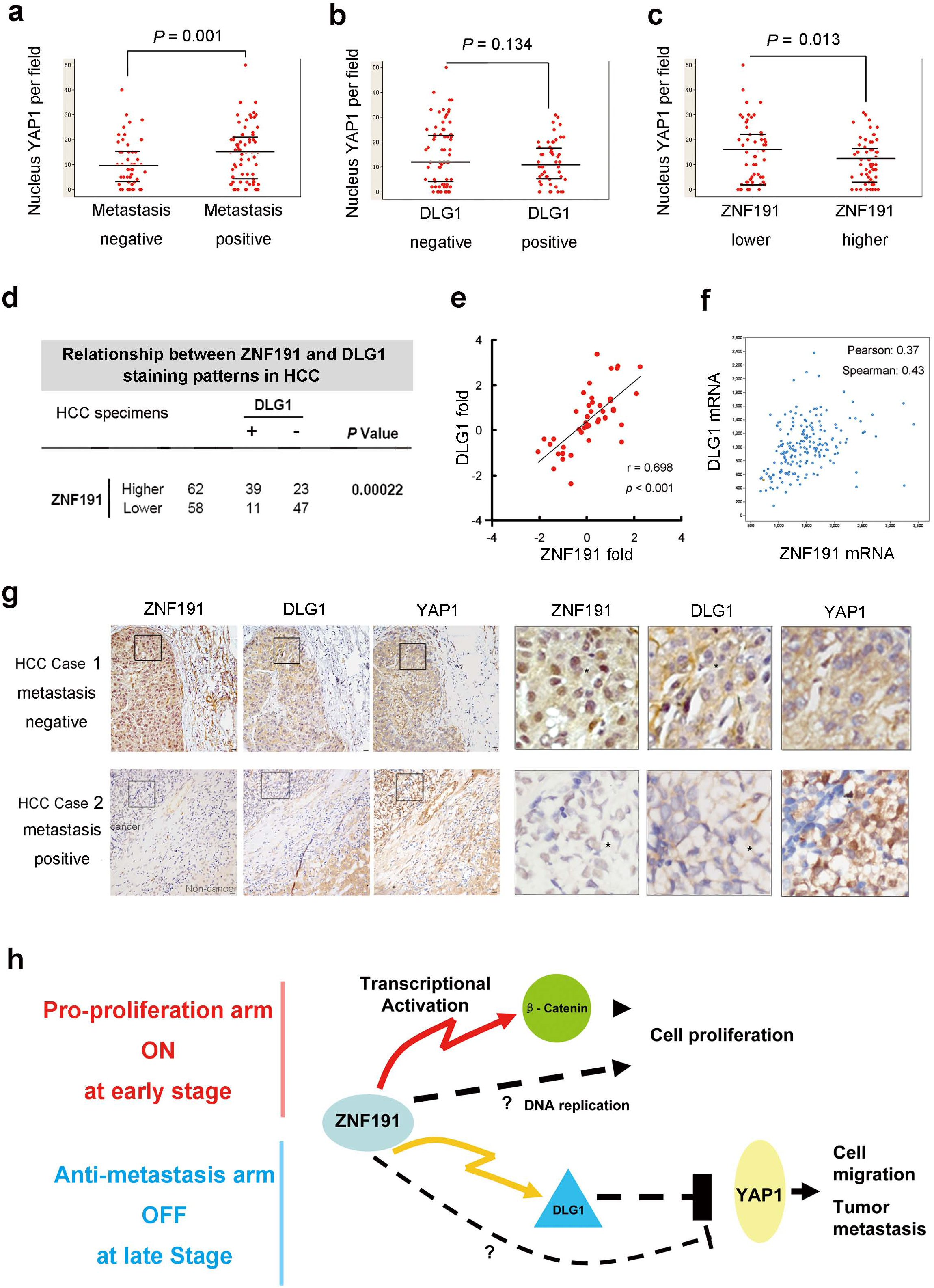
Inverse association between ZNF191 and nucleus YAP1, and the co-expression of ZNF191 and DLG1 in HCCs. (a) ∼ (d) Consecutive sections of 120 cases HCC tissue samples were subjected to IHC staining, for ZNF191, DLG1 and YAP1 respectively. (a) Also in Supplementary Fig. 10h. (b) DLG1 staining status in HCC specimens was correlated with YAP1 activation status. (c) ZNF191 staining status in HCC specimens was correlated with YAP1 activation status. (d) Relationship between ZNF191 and DLG1 staining status in HCC specimens. (e) In 44 human HCC tissues previous reported, ZNF191 and DLG1 mRNA fold changes compared to non-tumor adjacent tissues were measured using qPCR and 2^-ΔΔ^Ct method. Correlation analysis was performed with Pearson’s correlation test. (f) ZNF191 and DLG1 mRNA expression levels are positively correlated in HCC specimens. Data from 190 HCC specimens was downloaded from TCGA database: http://www.cbioportal.org/study.do?cancer_study_id=lihc_tcga, and analyzed at the cbioportal website. Unit of axis: RNA-sequencing V2 RSEM. Pearson’s or Spearman’s correlation coefficients were listed in the panels. (g) The typical ZNF191 staining was in the nucleus, but cytoplasm signal was also seen. The typical DLG1 staining was at membrane or in cytoplasm. YAP1 staining was typically in cytoplasm, sometimes in the nucleus at the rim of cancer foci. Two representative cases are shown. * Pinpoints as the reference points in consecutive sections. Bar = 25 μm. (h) The working model.

Although there are more DLG1 negative cases in YAP1 activating or metastasis cases, the correlation was neither significant statistically (Fig. 6b and Supplementary Table 2). DLG1 negative was actually correlated with high serum AFP levels and larger tumor size (Supplementary Table 2), indicating its tumor-suppressing function. This discrepancy implicates the existence of other mediator down-stream of ZNF191 and up-stream of YAP1.

## Discussion

Findings from our present study and others imply duel functions of ZNF191 in cancer (Fig. 6h). The model suggests that despite the pro-proliferation role, tumor cells would rather turn off ZNF191 expression in metastasis stage. Both the survival analysis in patents and *in vivo* experiments in mice confirmed that, although unfavorable for primary tumor growth, ZNF191 loss ultimately favors the dissemination of HCC cells to the whole body. Interestingly, DLG1, a *bona fide* target of ZNF191 identified in the current study, seems to be another ‘Jekyll and Hyde’ in tumor progression^23^. DLG1 accumulates in non-metastatic tumors but disappears in metastases^24^, sharing the same expression pattern as ZNF191 in HCC.

It was documented in *Drosophila* that *lgl*, *scrib* and *dlg* cooperated in a common genetic pathway^5^. *lgl* suppressed cell proliferation and survival via controlling *hippo*^9^; *scrib* knockdown caused activation of Yki (homolog of mammal YAP1)^8^; and *dlg* was found to suppress Yki in *Drosophila* follicle epithelial cells^11^and imaginal discs^8^. We showed that DLG1 knockdown in HCC cell lines HepG2 and MHCC-97H reduced YAP1 phosphorylation and blocked YAP1 from the regulation by ZNF191.

However, observations in HCC specimens failed to validate the association between DLG1 and YAP1 inactivation, but still confirmed the correlation between ZNF191 and YAP1 inactivation. These implicated alternative mechanisms that link ZNF191 and YAP1 signaling.

Screening binding factors of a critical element in MSLN promoter, ZNF191 was found co-localized with TEAD family members at an identical DNA motif^25^. This cryptic fact implies that ZNF191 may regulate YAP1 target genes through affecting TEAD. However, this hypothesis has not been validated because our ChIP-seq results failed to identify ZNF191-bound peaks in promoters of *CTGF*, *CYR61* or *EDN1*, where it should be occupied by TEADs^26^.

We demonstrated previously that ZNF191 activates the transcription of *CTNNB1* in HCC^12^. In the present study we found that ZNF191 inhibits YAP1 activation. Since `β`-catenin and YAP1 are generally redeemed over-lapped in functions and often co-existing^27^, our work has actually implied a paradox: If both regulations imposed by ZNF191 exist in HCC and functionally predominant, the activation of `β`-catenin and YAP1 should be mutually exclusive. Most intriguingly, a similar phenomenon, that nucleus`β`-catenin and nucleus YAP1 tend to exclude each other (over-lapped in less than 4% HCC cases), has been observed in HCC in a recent study^28^.

Our research suggests that ZNF191 positive and negative HCCs are distinct in clinical outcomes and driven by different mechanisms. To treat HCC precisely in the future, `β`-catenin or YAP1 signaling has to be targeted respectively according to the disease context including ZNF191 status. Restoring ZNF191 expression while limiting its pro-proliferation effects is a challenge, but offers a chance to block metastasis specifically. Therefore, we identify a therapeutic target that could be harnessed to prevent HCC metastasis and favor patient survival.

**Figure Legends** (Wu et al. ZNF191 Inhibits Cell Migration and Metastasis of Hepatocellular Carcinoma by DLG1-mediated YAP1 Inactivation)

**Supplementary Information** (Wu et al. ZNF191 Inhibits Cell Migration and Metastasis of Hepatocellular Carcinoma by DLG1-mediated YAP1 Inactivation)

**This Word file includes:**

Supplementary Discussion

Supplementary Figures Legends 1-10

Supplementary References

Caption for Supplementary Tables 3-8 (see separate Excel file for Supplementary Tables 3-8)

## Supplementary Discussion

### ZNF191 regulates YAP1 target genes in opposite directions in HEK293 cells

In HEK293 cells, there seem to be a paradox that ZNF191 should regulate YAP1 target genes in opposite directions. When we knock-down ZNF191 transiently, as manipulated in the previous transcript profiling^1^, the CTGF mRNA will reduce (Supplementary Fig. 7b); but if we stably deplete ZNF191 using lentivirus, CTGF and other YAP1 target genes will increase expression (Supplementary Fig. 7c).

We have noticed that DLG shares a similar regulation pattern towards YAP target genes. In Drosophila, when *dlg* (homologue of mammal *DLG*) was knocked-down moderately, *Yki* (homologue of *YAP*) target gene *diap1* should decrease. Only when *dlg* were depleted to the level where cells had lost polarity, would the *Yki* target genes be up-regulated^2^. Since ZNF191 regulates DLG1, that ZNF191 could mirror this regulation pattern. But the detailed mechanisms must be clarified in further investigation.

### CTGF is the marker of YAP1 activation, but unlikely the mediator of ZNF191’s function

In HCC cells, ZNF191 inhibits YAP1 target genes including CTGF. But CTGF seems not mediate of the function of ZNF191, because: (a) CTGF protein levels were scarcely affected after ZNF191 knockdown in HepG2 cells (data not shown); (b) we found CTGF had minor or even inhibitory effects upon HepG2 migration (Supplementary Fig. 6d, e); (c) Verteporfin disrupts YAP-TEAD^3^ and inhibits YAP1 target genes including CTGF^4^ but did not interfere with ZNF191’s migration-suppressing potential (Fig. 3e). It seemed that ZNF191 inhibits cell migration by inactivating YAP1 but not CTGF. TAZ (also known as WWTR1) is functionally paralleled with YAP1 and promotes epithelial-mesenchymal transition^5^. Selecting YAP1 as representative of YAP/TAZ signaling, does not necessarily mean YAP1 and TAZ are regulated equally by ZNF191, since ZNF191 has not shown its influence on epithelial-mesenchymal transition. Further study is still needed to investigate to what extent that TAZ could be affected by ZNF191-DLG1 signaling.

### The up-stream regulators of ZNF191

The regulators of ZNF191 remain elusive. We have noticed a very recent study, in searching for down-stream effectors of microRNA-940, ZNF191 was again found to be a suppressor of metastasis in gastric cancer with unclear mechanism^6^. This work not only supports our first finding (ZNF191 suppresses HCC metastasis) back to back, indicating that ZNF191 may be a general metastasis suppressor in a wider range of tumors, but also shed light onto the mechanisms controlling ZNF191 expression. We are interested to know whether microRNA-940 regulates ZNF191 in HCC. Considering the complex network of microRNA regulation, ZNF191 could be inhibited by means other than microRNA-940, finding regulators of ZNF191 at levels of transcription, pre-translation and post-translation is an inviting gap for future investigations.

## Supplementary Figure Legends

**Supplementary Figure 1.**
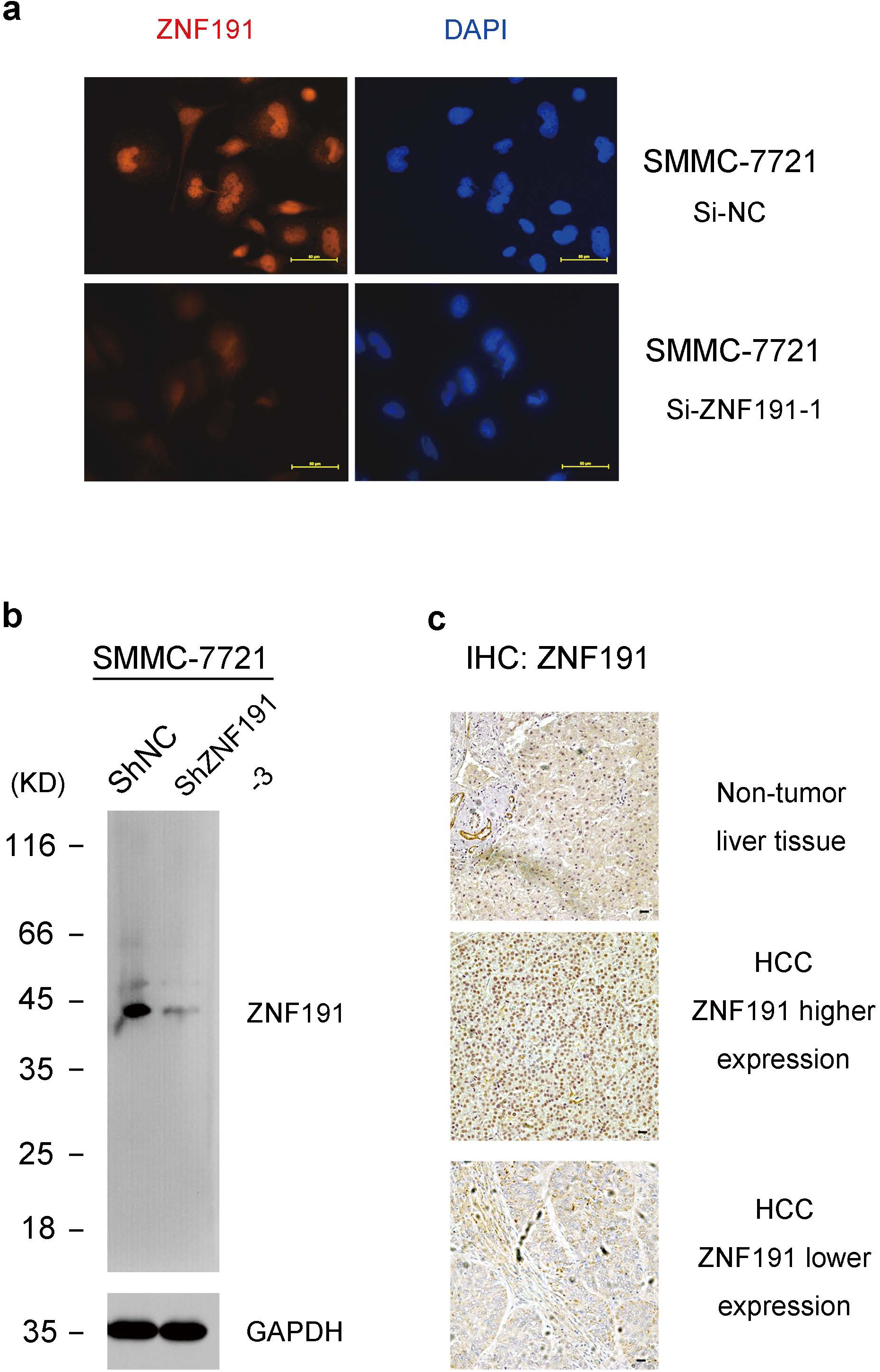
Validating the specificity of ZNF191 antibody for Immunohistochemistry (IHC). (a) Fixed cells were incubated with rabbit polyclonal anti-ZNF191 antibody (from SIGMA, HPA024062) and detected by immunofluorescence. The effect of RNAi is validated in Supplementary Fig. 4b. After depleting ZNF191 using RNAi, the fluorescence signal in the nuclei almost disappeared, leaving weak cytoplasm signal. Bar = 50 μm. (b) The same antibody as in (a) was applied to detect ZNF191 in cell lysates using Western Blotting. (c) Sections of HCC samples were incubated with the same ZNF191 antibody and stained by IHC. Representative images of higher or lower ZNF191 staining in HCC tissues were shown. Bar = 25 μm.

**Supplementary Figure 2.**
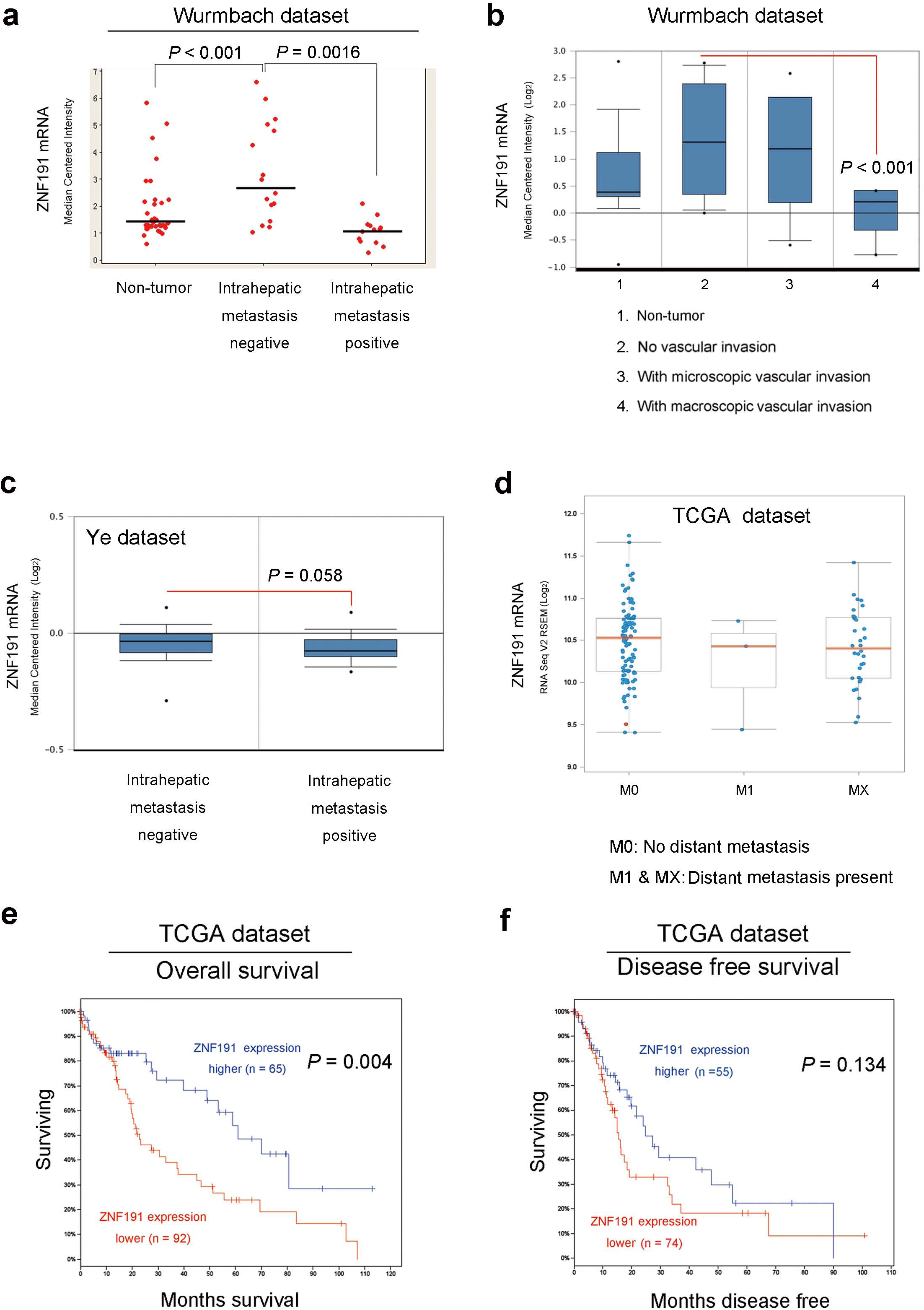
A twist of ZNF191 mRNA expression in metastatic HCC. (a) ZNF191 mRNA levels in metastasis-negative/positive or (b) in vascular-invasion-negative/positive HCCs (HCV-induced) compared with non-tumor samples were detected using microarray hybridization. Data were collected from Oncomine database (https://www.oncomine.org). The original data can be retrieved in reference^7^. Bar: median value. (c) In another dataset^8^, ZNF191 mRNA levels in metastasis-negative/positive HCCs (HBV positive) were revealed by Oncomine data-mining. (d) ZNF191 mRNA levels in 190 patients at different metastasis stages of HCC were demonstrated via RNA-sequencing. (e) & (f) Kaplan-Meier analyses of the correlations between ZNF191 mRNA levels and overall survival (e) or disease-free survival (f) of 190 patients with HCC. The mean expression level was used as the cutoff. Patients lacking surviving data were excluded. For (d) - (f), Data were collected and analyzed at the website: http://www.cbioportal.org, the original data can be retrieved from The Cancer Genome Atlas (TCGA) database: http://www.cbioportal.org/study.do?cancer_study_id=lihc_tcga. Publication permission: http://cancergenome.nih.gov/publications/publicationguidelines.

**Supplementary figure 3.**
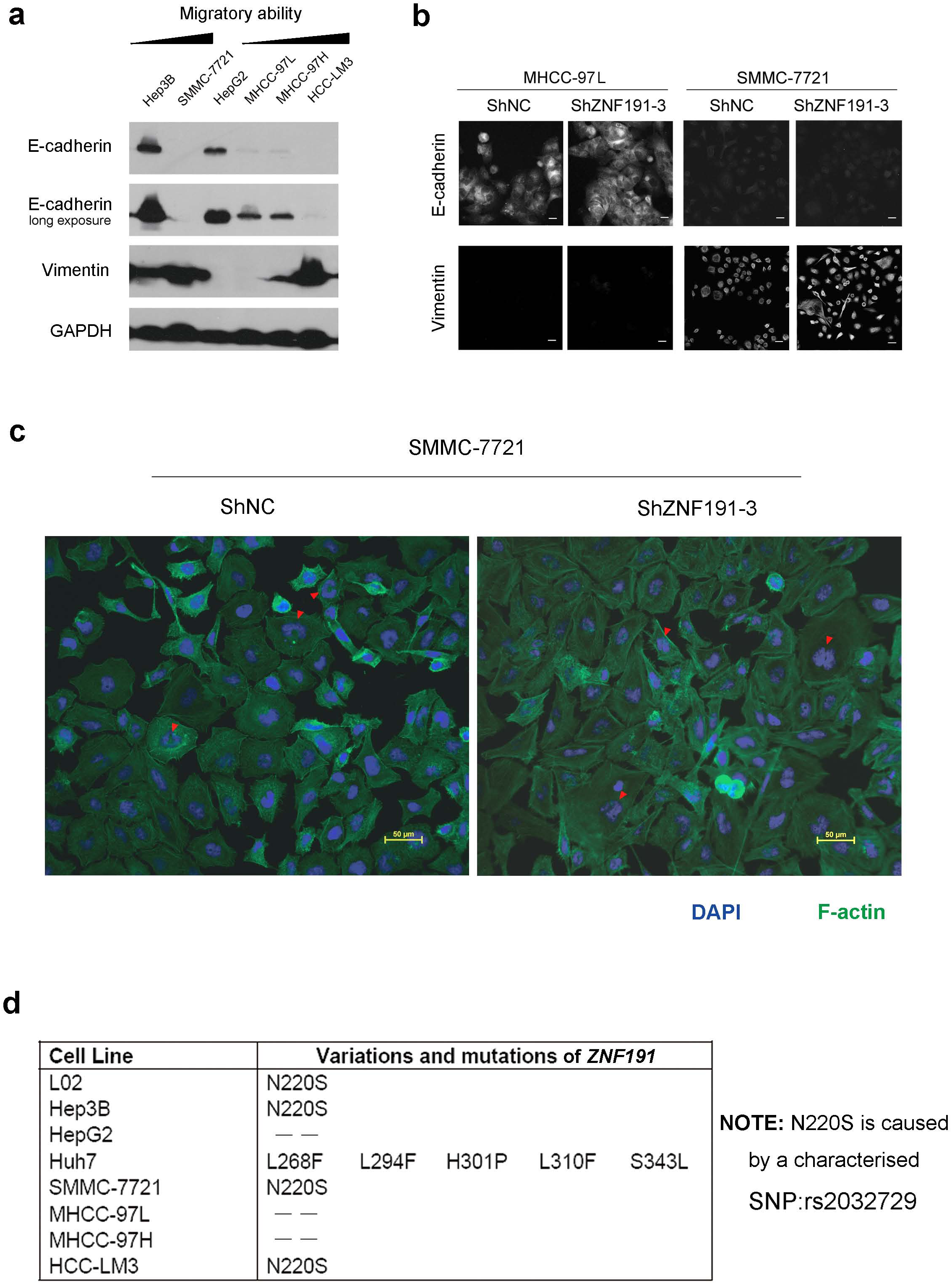
Identification of cell lines. (a) Equal amount of cell lysates were subjected to Western blotting. GAPDH was detected as the loading control. (b) Cells were subjected to immunofluorescence. Bar = 25 μm. (c) F-actin was dyed with fluorescence-labeled phalloidin. Red arrows designate the double-nuclei. (d) ZNF191 cDNA was cloned from each cell line and sequenced. NM_006965.3 was used as the reference mRNA of human ZNF191.

**Supplementary figure 4.**
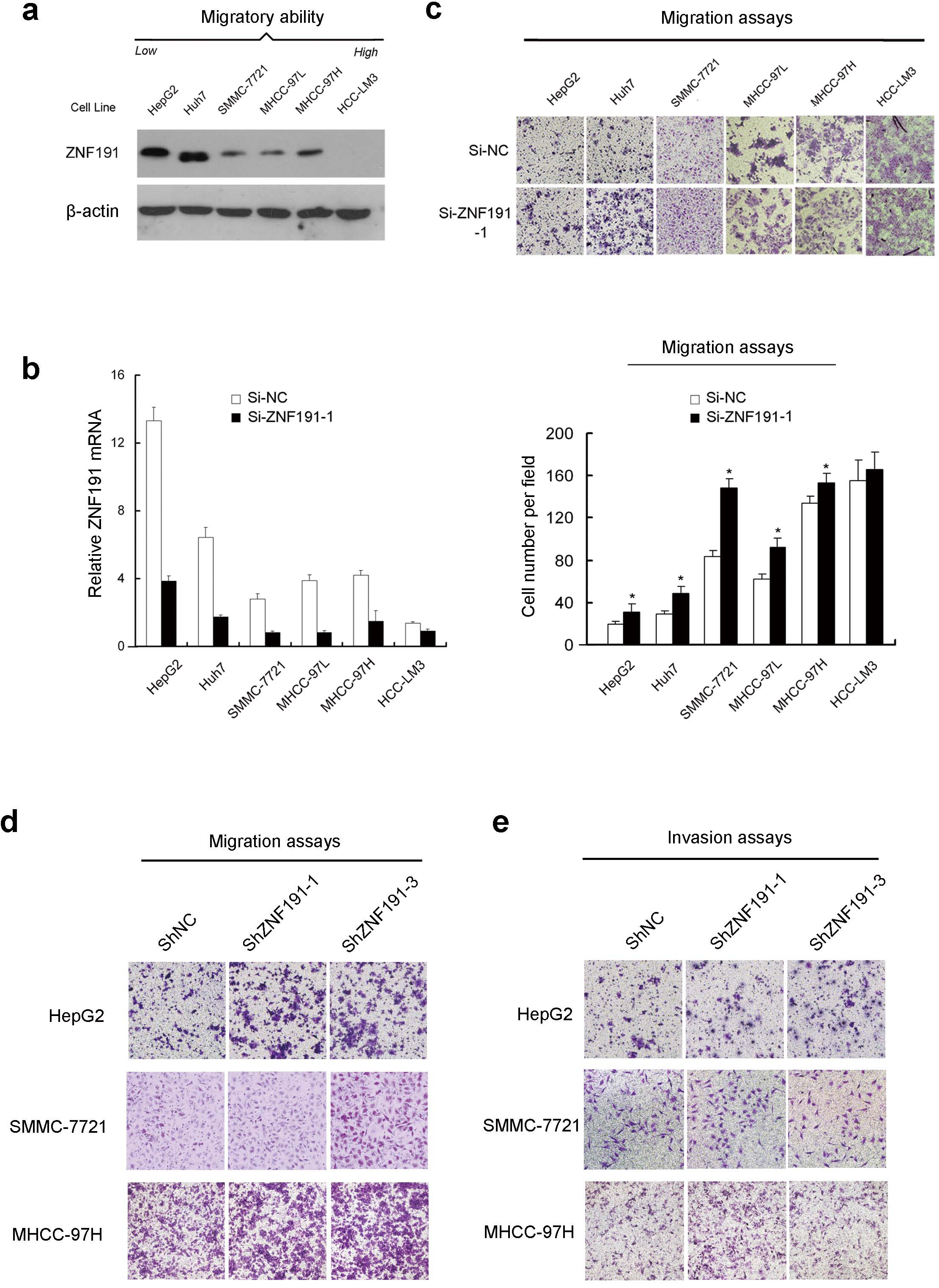
Functional effects of ZNF191 on HCC cell migration and proliferation *in vitro*. (a) The relative expression of ZNF191 in migratory HCC cell lines was detected by Western blotting. (b) The relative mRNA levels of ZNF191 in HCC cell lines and the efficacy of RNA interference (RNAi) were confirmed by quantitative real-time PCR (qPCR) and 2^-ΔΔ^Ct method. (c) HCC cells were transfected with indicated small interfering RNAs (siRNAs) for 48 hours. An equal number of the cells (50,000) were submitted to migration assays. (d) & (e) Representative results of Transwell based (d) migration and (e) invasion assays of different HCC stable lines. * *P* < 0.05.

**Supplementary figure 5.**
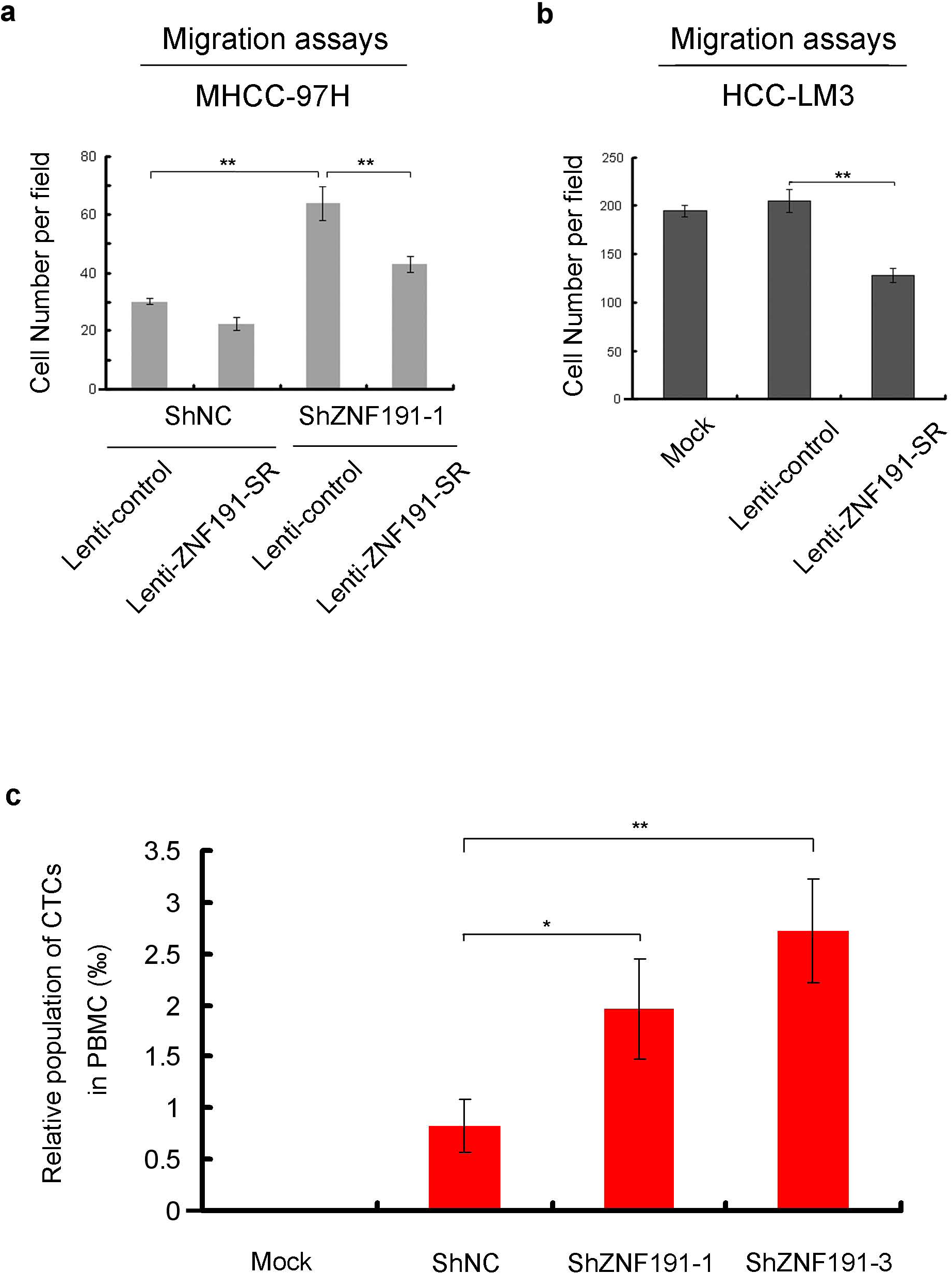
Statistic analysis of the results in Fig. 1 & 2. (a) & (b) Statistic analysis of the results described in Fig. 1f. (a) Over-expressing or rescuing ZNF191 in MHCC-97H cells. (b) Over-expressing ZNF191 in HCC-LM3 cells. (c) Statistic analysis of the results described in Fig. 2e. PBMC: Peripheral blood mononuclear cells. * *P* < 0.05; ** *P* < 0.01.

**Supplementary Figure 6.**
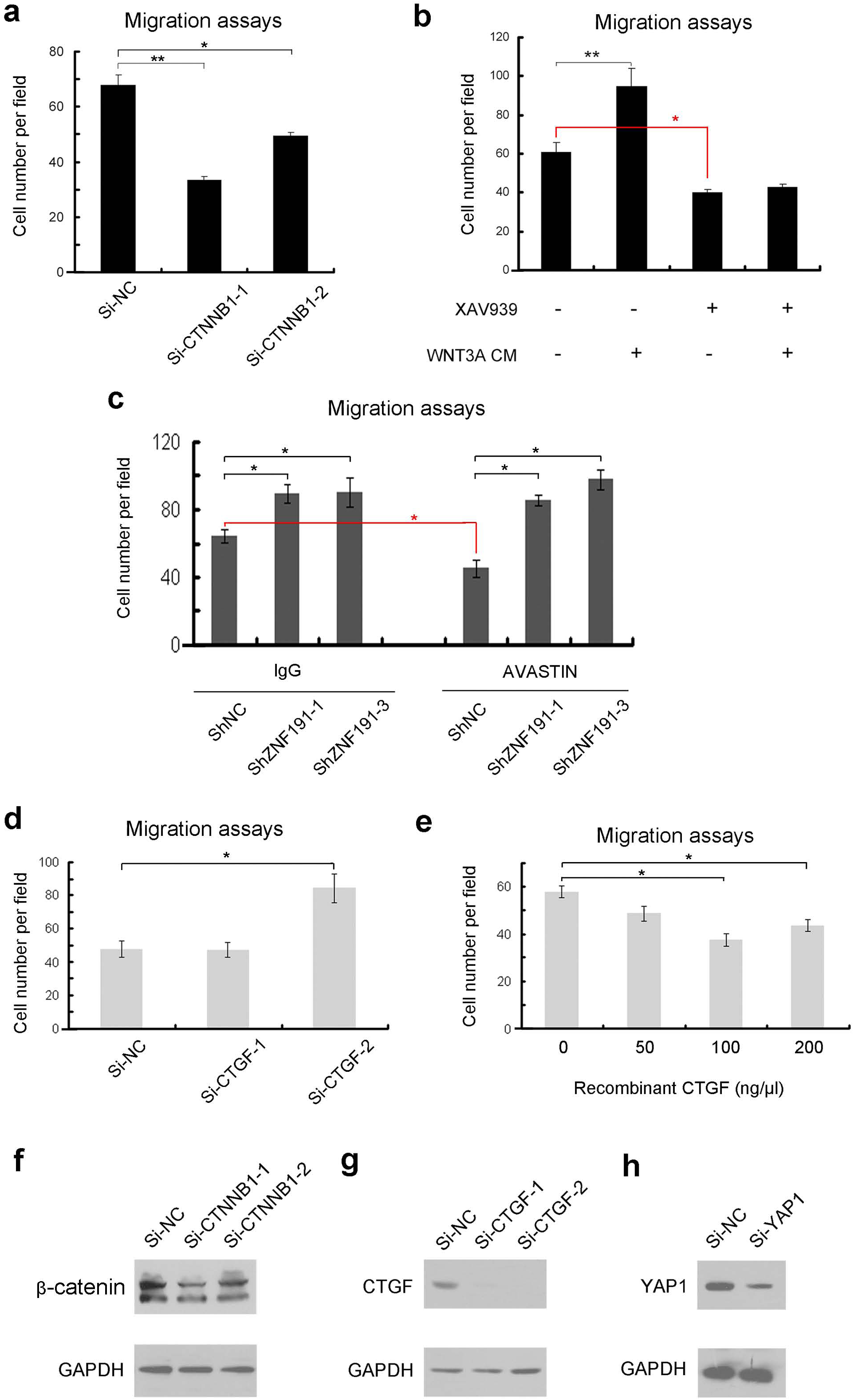
WNT/`β`-Catenin signaling, VEGF or CTGF does not mediate ZNF191’s function in cell migration. (a) 48 hours after siRNA transfection, HepG2 cells were subjected to migration assays. (b) HepG2 cells were treated with XAV939 (an inhibitor of WNT/`β`-Catenin) at 5 μM (DMSO as negative control) for 24 hours before assayed for migration ability. WNT3A conditional medium (CM, medium with 10% FBS as normal control) was used as attractant. (c) HepG2 stable lines were treated with VEGF-A-inhibiting antibody (AVASTIN) or normal human IgG at 10 ng/μl during the migration assays. (d) 48 hours after transfection with indicated siRNAs, HepG2 cells were subjected to migration assays. (e) HepG2 cells were treated with CTGF protein at gradient concentrations during the migration assays. (f) - (h) To minimize side effects, non-liposome transfection reagent INTERFERin (see Supplementary Materials & Methods) was applied and the final concentration of siRNA was 2.5nM for each transfection. The knockdown of each protein was validated by Western blotting. The two-band blotting of `β`-CATENIN is detailed in **Cell line identification and choosing** section of **Online Methods**. * *P* < 0.05; ** *P* < 0.01.

**Supplementary Figure 7.**
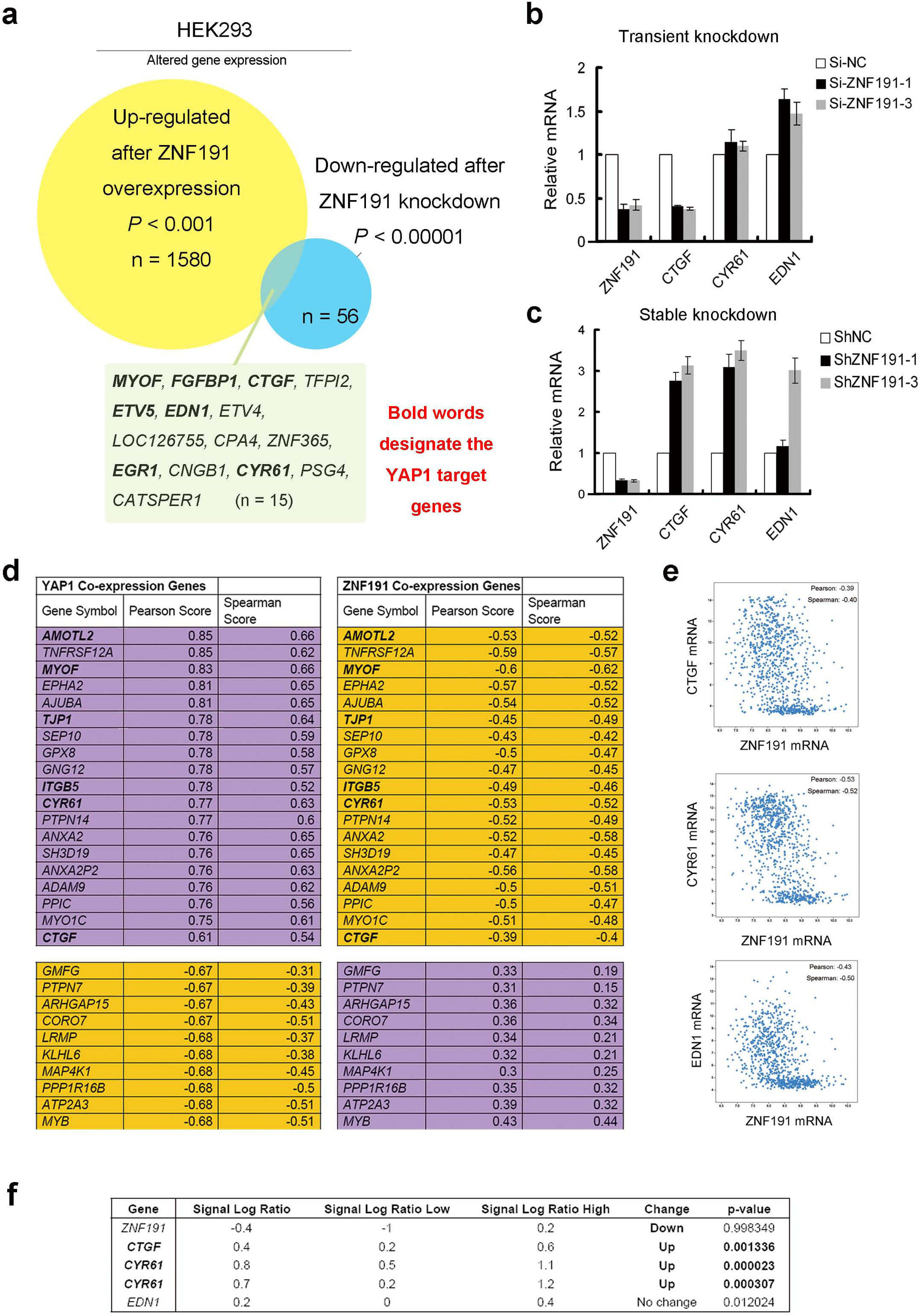
ZNF191 is involved in YAP1 signaling. (a) Re-analysis of the transcript profiling data from a precious work^1^. The overlap region in the Venn diagram represents the set of genes regulated by ZNF191 most significantly in HEK293 cells. Nearly half of them (7 in 15) are known YAP1 target genes ^9, 10^. (b) HEK293 cells were transfected with indicated siRNAs for 24 hours before extracted of RNA. Fold changes of mRNA levels were quantified with qPCR. (c) ZNF191 was knocked-down stably in HEK293 cells through lentivirus mediated shRNA expression. Fold changes of mRNA levels were quantified with qPCR. (d) Gene expression profiles in 883 lines of cancer cells defined the 20 most positively (colored purple, 19 genes shown excluding YAP1 itself) and the 10 most negatively (colored orange) co-expressed genes with YAP1, listed in the left column. For each of these genes, the correlation coefficient (as Pearson or Spearman Scores) with ZNF191 expression was listed in the right column. The bold words indicate the down-stream target genes of YAP1^9, 10^. (e) Scatter plots showed the reverse correlation of mRNA expression between ZNF191 and 3 well-defined YAP1 target genes *CTGF*, *CYR61* and *EDN1* in 883 cancer cell lines. For (d) & (e), the analysis was performed at the website: http://www.cbioportal.org. Original data can be retrieved from TCGA database: http://www.cbioportal.org/study.do?cancer_study_id=cellline_ccle_broad, or from the reference ^11^. Unit of axis: RNA-sequencing V2 RSEM. Pearson’s or Spearman’s correlation coefficients were listed in the panels. (f) L02 cells were stably transfected with negative control or ZNF191-depleting shRNA; the mRNA levels were profiled with microarrays. Shown in the figure is part of our data in a previous work^12^.

**Supplementary Figure 8.**
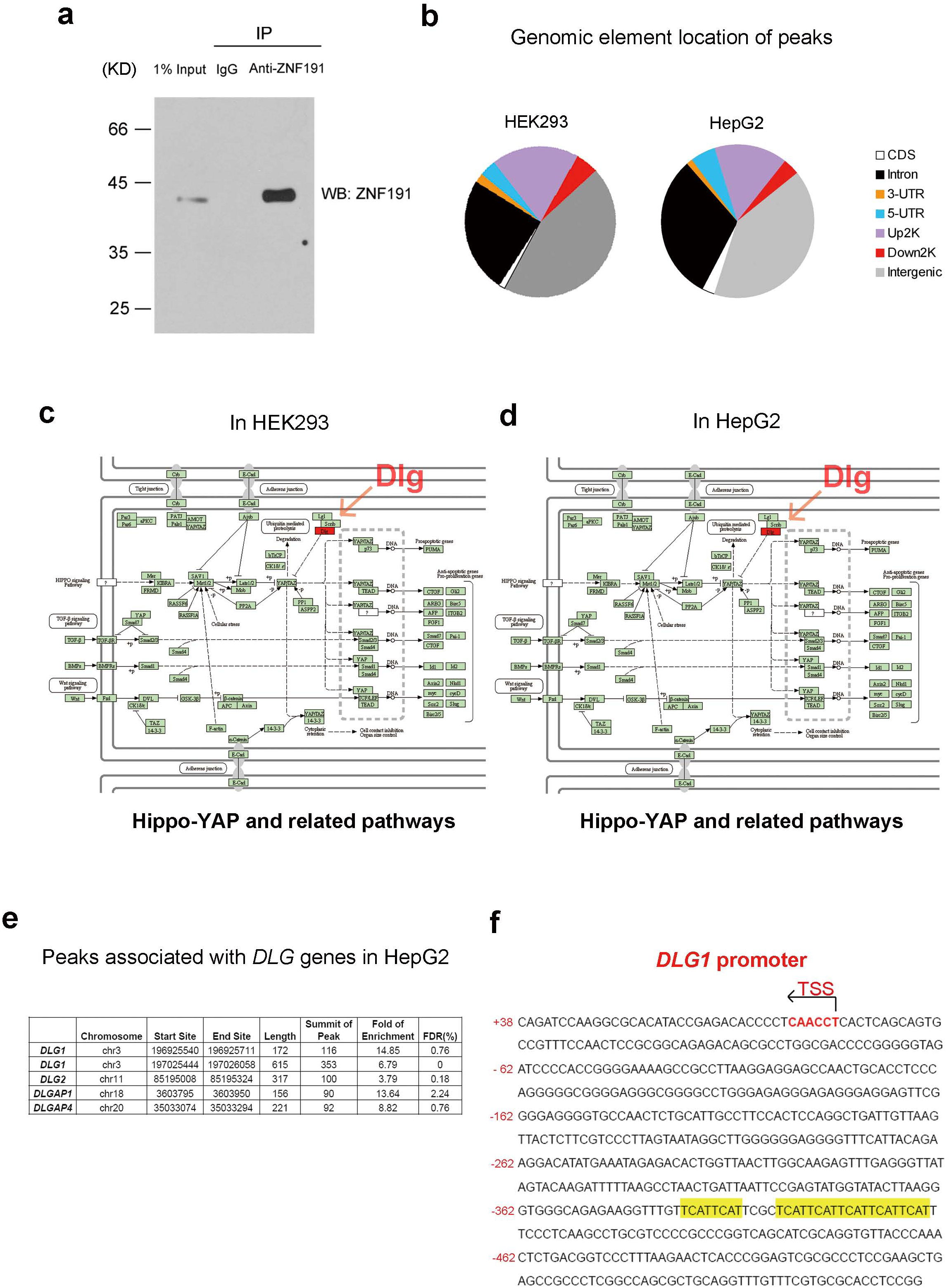
*DLG1* is the only YAP signaling regulator with its promoter bound by ZNF191. (a) Validating the specificity and affinity of the rabbit polyclonal anti-ZNF191 antibody (from Abcam, ab176589) for immunoprecipitation in HEK293 cells. (b) Statistic analysis of the peaks shows the enrichment of ZNF191-bound peaks in genome elements. (c) & (d) Pathway analysis was performed using the KEGG database. Gene with its promoter (upstream 2000bp region of TSS) bound by ZNF191 in HEK293(c) and HepG2 (d) cells would be colored red in the schematic pathway diagram. (e) In a set of genes with their whole regions (upstream 2000bp region of TSS, gene body and downstream 2000bp of TTS) bound by ZNF191, we found more *DLG* gene family members. (f) The sequence of *DLG1* proximal promoter. Highlighted are the TCAT repeating motifs.

**Supplementary Figure 9.**
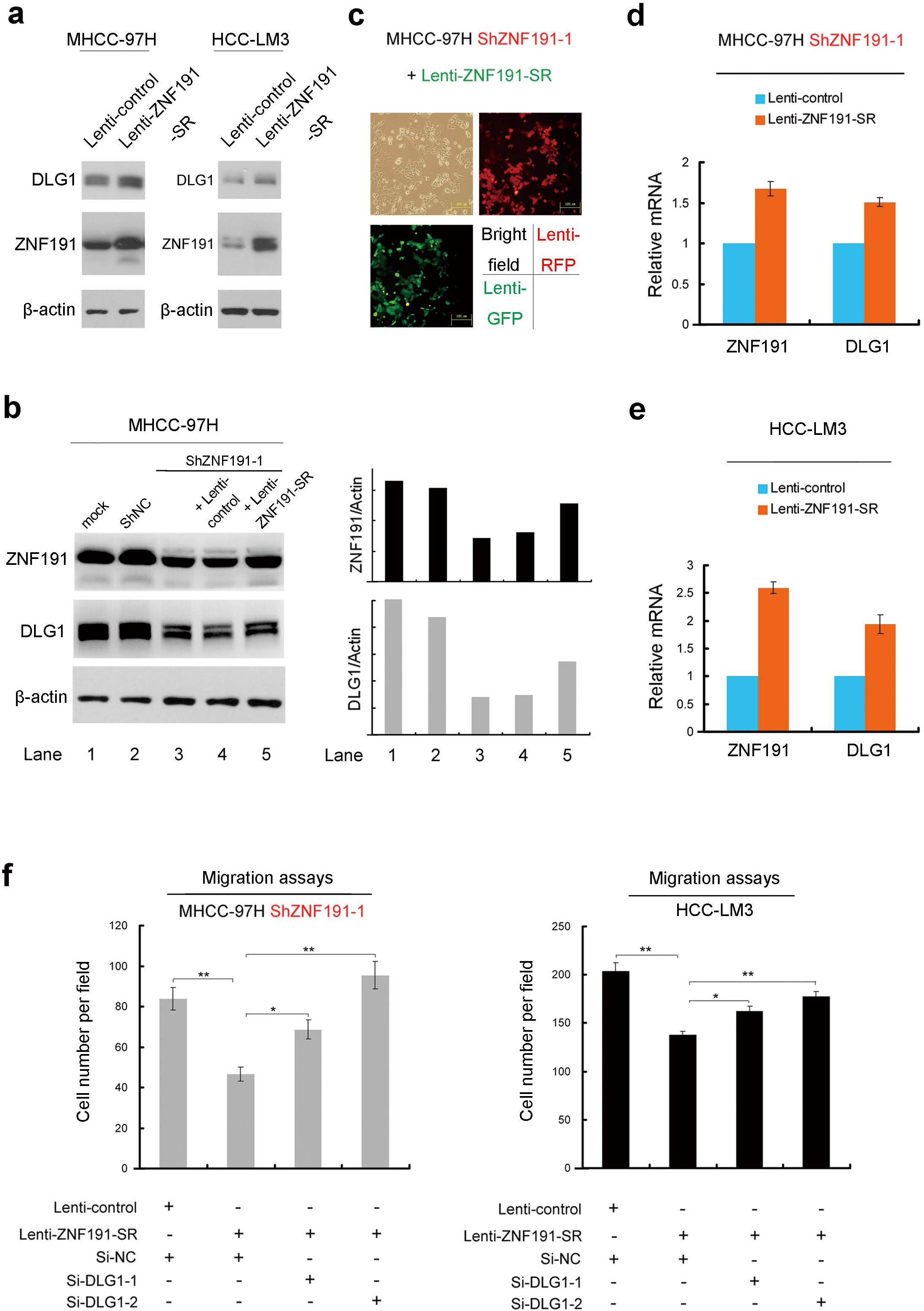
Over-expressing or rescuing ZNF191 increased DLG1 expression levels, inhibited cell migration. (a) & (b) Cells transfected with lentivirus had been selected and maintained for 20 days. Equal amounts of cell lysates were subjected to Western blotting. Grayscale scanning and analysis were performed as in Figure 3b. (c) Lentivirus expressing ShZNF191-1 co-expressed RFP, while lentivirus expressing ZNF191 cDNA co-expressed EGFP. Rescuing percentage was validated by over-lapping the two kinds of fluorescence signal. (d) & (e) Fold changes of mRNA levels were quantified by qPCR. (f) Stable cell lines were transfected with indicated siRNAs for 48 hours and subjected to migration assays.

**Supplementary Figure 10.**
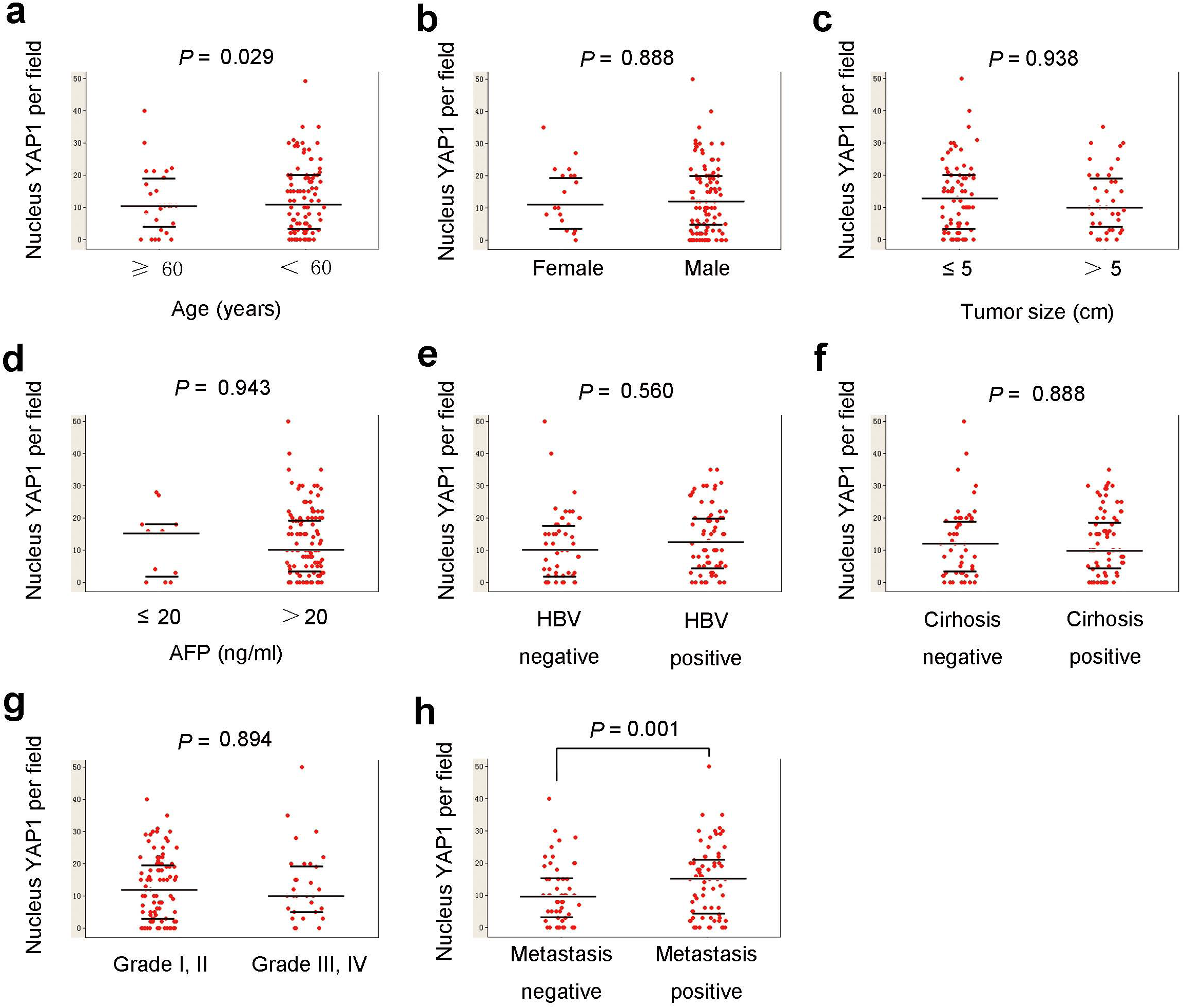
Correlation of YAP1 activation levels in HCC specimens with their clinicopathological features. (a) ∼ (h) For each sample, five random fields were selected and mean number of nucleus YAP1 in these fields was recorded as the YAP1 activation level in this sample. Grouped by different clinicopathological features as (a) age, (b) gender, (c) tumor diameter, (d) serum AFP level, (e) HBV infection status, (f) cirrhosis status, (g) Edmondson’s grade or (h) intrahepatic metastasis status, *P* value shows the significance of YAP1 activation level alternation between the two groups, as listed for each clinicopathological feature.

---

**Supplementary Table 3** Primers, Probes and RNAi target sequences

**Supporting Table 4** Antibody List

**Supporting Table 5** Compounds and Special reagent

**Supporting Table 6** ZNF191-bound genes in HEK293 (2000bp up-stream of TSS)

**Supporting Table 7** ZNF191-bound genes in HepG2 (2000bp up-stream of TSS)

**Supporting Table 8** Results of statistic analysis in HCC specimens

**Table 1.** Relationships between ZNF191 expression and clinicopathological features in HCC tissues

| Clinical Features | Cases | ZNF191 Staining (Cancer) |  | P Value |
| --- | --- | --- | --- | --- |
|  |  | Higher<br>(n = 62)<br>Cases (%) | Lower<br>(n = 58)<br>Cases (%) |  |
| **Age (years)** |  |  |  |  |
| <60 | 93 | 46(49.5) | 47(50.5) | 0.533 |
| ≥60 | 27 | 16(59.3) | 11(40.7) |  |
| **Gender** |  |  |  |  |
| Female | 18 | 9(50.0) | 9(50.0) | 0.915 |
| Male | 102 | 53(52.0) | 49(48.0) |  |
| **HBsAg** |  |  |  |  |
| Negative | 49 | 26(53.1) | 23(46.9) | 0.860 |
| Positive | 71 | 36(50.7) | 35(49.3) |  |
| **Liver Cirrhosis** |  |  |  |  |
| Negative | 51 | 23(45.1) | 28(54.9) | 0.389 |
| Positive | 69 | 39(56.5) | 30(43.5) |  |
| **Serum AFP (ng/ml)** |  |  |  |  |
| ≤20 | 11 | 6(54.5) | 5(45.5) | 0.889 |
| >20 | 109 | 56(51.4) | 53(48.6) |  |
| **Tumor Size (cm)** |  |  |  |  |
| ≤5 | 81 | 47(58.0) | 34(42.0) | 0.162 |
| >5 | 39 | 15(38.5) | 24(61.5) |  |
| **Edmondson's Grade** |  |  |  |  |
| I,II | 90 | 51(56.7) | 39(43.3) | 0.187 |
| III,IV | 30 | 11(36.7) | 19(63.3) |  |
| **Intrahepatic Metastasis** |  |  |  |  |
| Negative | 51 | 37(72.5) | 14(27.5) | **0.006** |
| Positive | 69 | 25(36.2) | 44(63.8) |  |

**Table 2.** Relationships between DLG1 expression and clinicopathological features in HCC tissues

| Clinical Features | Cases | DLG1 Staining (Cancer) |  | P Value |
| --- | --- | --- | --- | --- |
|  |  | Positive<br>(n = 50)<br>Cases (%) | Negative<br>(n =70)<br>Cases (%) |  |
| **Age (years)** |  |  |  |  |
| <60 | 93 | 36(38.7) | 57(61.3) | 0.352 |
| ≥60 | 27 | 14(51.8) | 13(48.2) |  |
| **Gender** |  |  |  |  |
| Female | 18 | 6(33.3) | 12(66.7) | 0.552 |
| Male | 102 | 44(43.1) | 58(56.9) |  |
| **HBsAg** |  |  |  |  |
| Negative | 49 | 25(51.0) | 24(49.0) | 0.188 |
| Positive | 71 | 25(35.2) | 46(64.8) |  |
| **Liver Cirrhosis** |  |  |  |  |
| Negative | 51 | 20(39.2) | 31(60.8) | 0.721 |
| Positive | 69 | 30(43.5) | 39(56.5) |  |
| **Serum AFP (ng/ml)** |  |  |  |  |
| ≤20 | 11 | 8(72.7) | 3(27.3) | **0.019** |
| >20 | 109 | 42(38.5) | 67(61.5) |  |
| **Tumor Size (cm)** |  |  |  |  |
| ≤5 | 81 | 41(50.6) | 40(49.4) | **0.029** |
| >5 | 39 | 9(23.1) | 30(76.9) |  |
| **Edmondson's Grade** |  |  |  |  |
| I,II | 90 | 38(42.2) | 52(57.8) | 0.870 |
| III,IV | 30 | 12(40.0) | 18(60.0) |  |
| **Intrahepatic Metastasis** |  |  |  |  |
| Negative | 51 | 26(51.0) | 25(49.0) | 0.176 |
| Positive | 69 | 24(34.8) | 45(65.2) |  |

**Footnote:**

Expression changes of ZNF191 (cancer vs. adjacent non-cancerous tissue) were analyzed by two eligible pathologists (discrepancies arbitrated by a third). P value represents the probability from a *χ* ^2^ test for different immunohistochemical staining of ZNF191 in HCC tissues.

**For boldfaced probability value:** *P* < 0.05.

**Intrahepatic Metastasis** means small HCC nodules such as venous permeation and tumor microsatellite nodules distributed around the primary (larger) HCC masses.

